# Tuning transcriptional regulation through signaling: A predictive theory of allosteric induction

**DOI:** 10.1101/111013

**Authors:** Manuel Razo-Mejia, Stephanie L. Barnes, Nathan M. Belliveau, Griffin Chure, Tal Einav, Mitchell Lewis, Rob Phillips

## Abstract

Allosteric regulation is found across all domains of life, yet we still lack simple, predictive theories that directly link the experimentally tunable parameters of a system to its input-output response. To that end, we present a general theory of allosteric transcriptional regulation using the Monod-Wyman-Changeux model. We rigorously test this model using the ubiquitous simple repression motif in bacteria by first predicting the behavior of strains that span a large range of repressor copy numbers and DNA binding strengths and then constructing and measuring their response. Our model not only accurately captures the induction profiles of these strains but also enables us to derive analytic expressions for key properties such as the dynamic range and [*EC*_50_]. Finally, we derive an expression for the free energy of allosteric repressors which enables us to collapse our experimental data onto a single master curve that captures the diverse phenomenology of the induction profiles.

## Introduction

Understanding how organisms sense and respond to changes in their environment has long been a central theme of biological inquiry. At the cellular level, this interaction is mediated by a diverse collection of molecular signaling pathways. A pervasive mechanism of signaling in these pathways is allosteric regulation, in which the binding of a ligand induces a conformational change in some target molecule, triggering a signaling cascade [1]. One of the most important examples of such signaling is offered by transcriptional regulation, where a transcription factor’s propensity to bind to DNA will be altered upon binding to an allosteric effector.

Despite the overarching importance of this mode of signaling, a quantitative understanding of the molecular interactions between extracellular inputs and gene expression remains poorly explored. Attempts to reconcile theoretical models and experiments have often been focused on fitting data retrospectively after experiments have been conducted [2, 3]. Further, many treatments of induction are strictly phenomenological, electing to treat induction curves individually either using Hill functions or as ratios of polynomials without acknowledging that allosteric proteins have distinct conformational states depending upon whether an effector molecule is bound to them or not [4–8]. These fits are made in experimental conditions in which there is great uncertainty about the copy number of both the transcription factor and the regulated locus, meaning that the underlying minimal set of parameters cannot be pinned down unequivocally. This leaves little prospect for predicting or understanding what molecular properties determine key phenotypic parameters such as leakiness, dynamic range, [*EC*_50_], and the effective Hill coefficient as discussed in Refs. [9, 10] and illustrated in Fig. 1. Our goal was to use a minimal Monod-Wyman-Changeux (MWC) model of transcription factor induction in conjunction with a corresponding thermodynamic model of repression to test whether such a simple model is capable of predicting how the induction process changes over broad swathes of regulatory parameter space. While some treatments of induction have used MWC models to predict transcriptional outputs [3, 11, 12], these often require multi-parameter fitting which gives rise to issues of parameter degeneracy (see Appendix A) and may include effective parameters that have tenuous biological meaning. In contrast, our objective was to use the MWC model to make parameter-free predictions about how the induction response will be altered when transcription factor copy number and operator strength are systematically varied.

**Figure 1.**
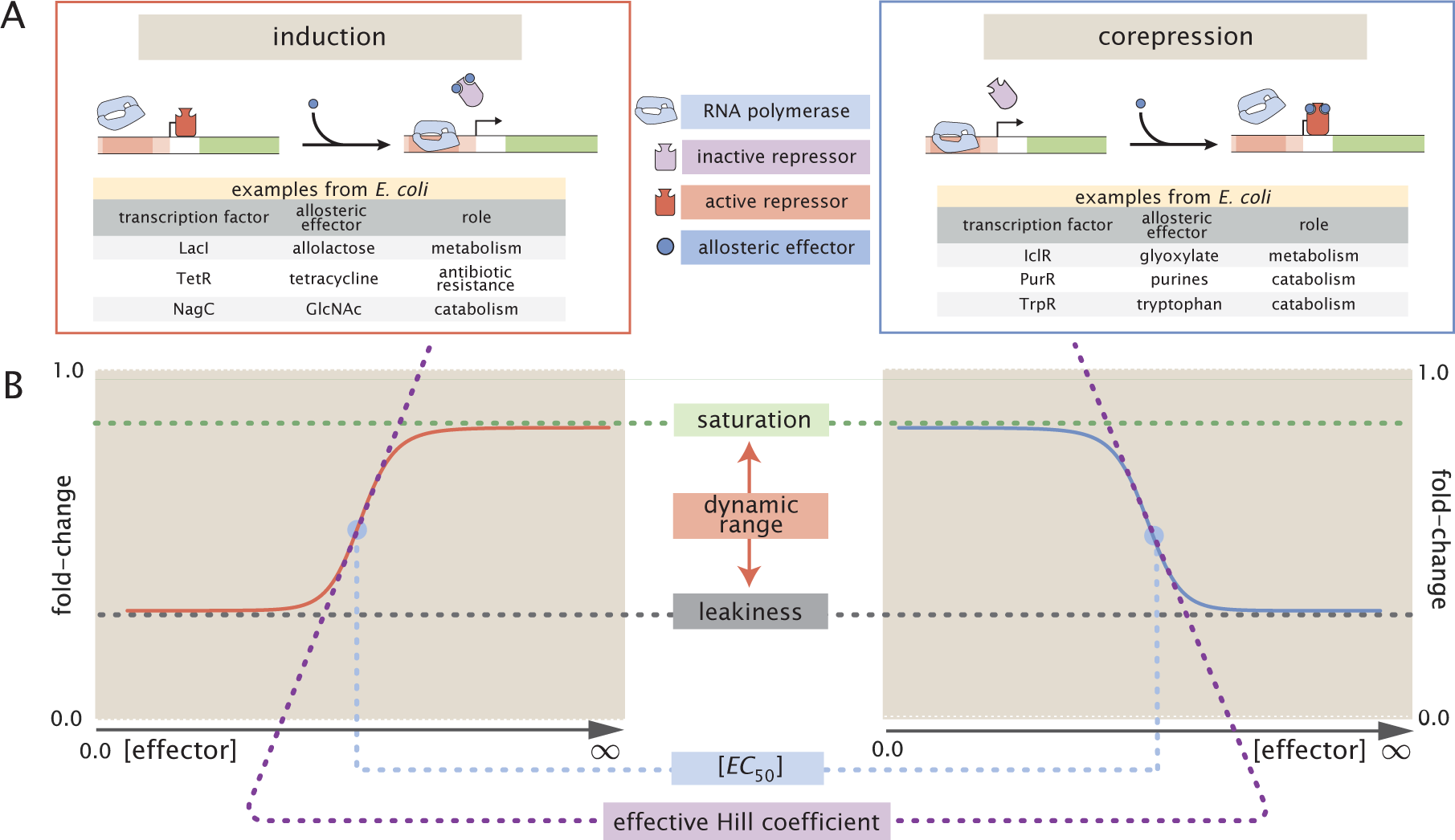
Transcription regulation architectures involving an allosteric repressor. (A) We consider a promoter regulated solely by an allosteric repressor. When bound, the repressor prevents RNAP from binding and initiating transcription. Induction is characterized by the addition of an effector which binds to the repressor and stabilizes the inactive state (defined as the state which has a low affinity for DNA), thereby increasing gene expression. In corepression, the effector stabilizes the repressor’s active state and thus further reduces gene expression. We list several characterized examples of induction and corepression that support different physiological roles in *E. coli* [25, 26]. (B) A schematic regulatory response of the two architectures shown in Panel A plotting the fold-change in gene expression as a function of effector concentration, where fold-change is defined as the ratio of gene expression in the presence versus the absence of repressor. We consider the following key phenotypic properties that describe each response curve: the minimum response (leakiness), the maximum response (saturation), the difference between the maximum and minimum response (dynamic range), the concentration of ligand which generates a fold-change halfway between the minimal and maximal response ([*EC*_50_]), and the log-log slope at the midpoint of the response (effective Hill coefficient).

We test our model in the context of the simple repression motif – a widespread bacterial genetic regulatory architecture in which binding of a transcription factor occludes binding of an RNA polymerase thereby inhibiting transcription initiation. A recent survey of different regulatory architectures within the *E. coli* genome revealed that more than 100 genes are characterized by the simple repression motif, making it a common and physiologically relevant architecture [13]. Building upon previous work [14–16], we present a statistical mechanical rendering of allostery in the context of induction and corepression, shown schematically in Fig. 1A, and use this model as the basis of parameter-free predictions which we then probe experimentally. Specifically, we model the allosteric response of transcriptional repressors using the MWC model, which stipulates that an allosteric protein fluctuates between two distinct conformations ‑ an active and inactive state – in thermodynamic equilibrium [17]. In the context of induction, effector binding increases the probability that a repressor will be in the inactive state, weakening its ability to bind to the promoter and resulting in increased expression. The framework presented here provides considerable insight beyond that of simply fitting a sigmoidal curve to inducer titration data. We aim to explain and predict the relevant biologically important parameters of an induction profile, such as characterizing the midpoint and steepness of its response as well as the limits of minimum and maximum expression as shown in Fig. 1B. By combining this MWC treatment of induction with a thermodynamic model of transcriptional regulation (Fig. 2), we create a general quantitative model of allosteric transcriptional regulation that is applicable to a wide range of regulatory architectures such as activation, corepression, and various combinations thereof, extending our quantitative understanding of these schemes [18] to include signaling.

**Figure 2.**
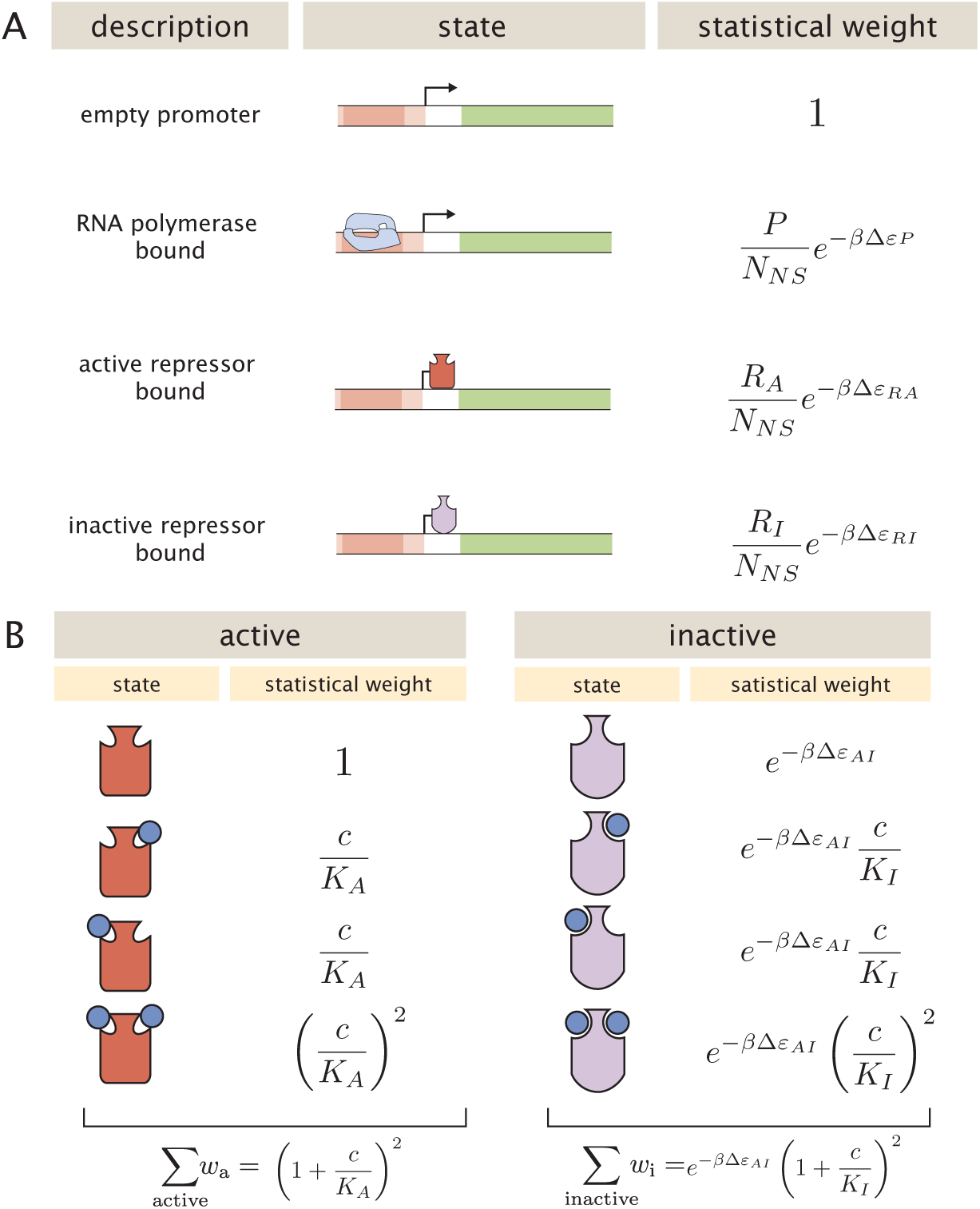
States and weights for the simple repression motif. (A) RNAP (light blue) and a repressor compete for binding to a promoter of interest. There are *R_A_* repressors in the active state (red) and *R_I_* repressors in the inactive state (purple). The difference in energy between a repressor bound to the promoter of interest versus another non-specific site elsewhere on the DNA equals Δ*ε_RA_* in the active state and Δ*ε_RI_* in the inactive state; the *P* RNAP have a corresponding energy difference Δ*ε_P_* relative to non-specific binding on the DNA. *N_NS_* represents the number of non-specific binding sites for both RNAP and repressor. (B) A repressor has an active conformation (red, left column) and an inactive conformation (purple, right column), with the energy difference between these two states given by Δ*ε_AI_*. The inducer (blue circle) at concentration *c* is capable of binding to the repressor with dissociation constants *K_A_* in the active state and *K_I_* in the inactive state. The eight states for a dimer with *n* = 2 inducer binding sites are shown along with the sums of the active and inactive states.

To demonstrate the predictive power of our theoretical formulation across a wide range of both operator strengths and repressor copy numbers, we design an *E. coli* genetic construct in which the binding probability of a repressor regulates gene expression of a fluorescent reporter. Using components from the well-characterized *lac* system in *E. coli*, we first quantify the three parameters associated with the induction of the repressor, namely, the binding affinity of the active and inactive repressor to the inducer and the free energy difference between the active and inactive repressor states. We determine these parameters by fitting to measurements of the fold-change in gene expression as a function of inducer concentration for a circuit with known repressor copy number and repressor-operator binding energy. We note that all other parameters that appear in the thermodynamic model are used without change from a suite of earlier experiments which quantify fold-change in a range of regulatory scenarios [14, 15, 19–22]. With these estimated allosteric parameters in hand, we make accurate, parameter-free predictions of the induction response for many other combinations of repressor copy number and binding energy. This goes well beyond previous treatments of the induction phenomenon and shows that one extremely compact set of parameters can be applied self-consistently and predictively to vastly different regulatory situations including simple repression on chromosome, cases in which decoy binding sites for repressor are put on plasmids, cases in which multiple genes compete for the same regulatory machinery, cases involving multiple binding sites for repressor leading to DNA looping, and the induction experiments described here. The broad reach of this minimal parameter set is highlighted in Fig. 3.

**Figure 3.**
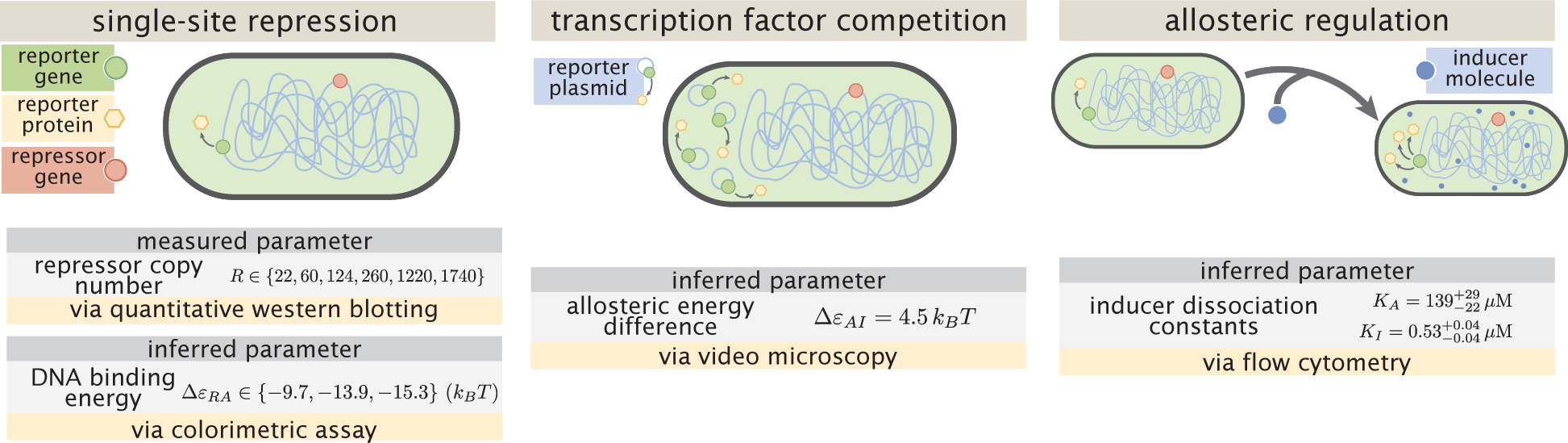
Understanding the modular components of induction. Over the past decade, we have refined both our experimental control over and theoretical understanding of the simple repression architectures. A first round of experiments used colorimetric assays and quantitative Western blots to investigate how single-site repression is modified by the repressor copy number and repressor-DNA binding energy [14]. A second round of experiments used video microscopy to probe how the copy number of the promoter and presence of competing repressor binding sites affect gene expression, and we use this data set to determine the free energy difference between the repressor’s inactive and active conformations [15] (see Appendix A). Both of the previous experiments characterized the system in the absence of an inducer, and in the present work we consider this additional important feature of the simple repression architecture. We used flow cytometry to determine the inducer-repressor dissociation constants and demonstrate that with these parameters we can predict *a priori* the behavior of the system for any repressor copy number, DNA binding energy, gene copy number, and inducer concentration.

Rather than viewing the behavior of each circuit as giving rise to its own unique input-output response, the formulation of the MWC model presented here provides a means to characterize these seemingly diverse behaviors using a single unified framework governed by a small set of parameters, applicable even to mutant repressors in much the same way that earlier work showed how mutants in quorum sensing and chemotaxis receptors could be understood within a minimal MWC-based model [23, 24]. Another insight that emerges from our theoretical treatment is how a subset in the parameter space of repressor copy number, operator binding site strength, and inducer concentration can all yield the same level of gene expression. Our application of the MWC model allows us to understand these degeneracies in parameter space through an expression for the free energy of repressor binding, a nonlinear combination of physical parameters which determines the system’s mean response and is the fundamental quantity that dictates the phenotypic cellular response to a signal.

## Results

### Characterizing Transcription Factor Induction using the Monod-Wyman-Changeux (MWC) Model

We begin by considering the induction of a simple repression genetic architecture, in which the binding of a transcriptional repressor occludes the binding of RNA polymerase (RNAP) to the DNA [27, 28]. When an effector (hereafter referred to as an “inducer” for the case of induction) binds to the repressor, it shifts the repressor’s allosteric equilibrium towards the inactive state as specified by the MWC model [17]. This causes the repressor to bind more weakly to the operator, which increases gene expression. Simple repression motifs in the absence of inducer have been previously characterized by an equilibrium model where the probability of each state of repressor and RNAP promoter occupancy is dictated by the Boltzmann distribution [14, 15, 27–30] (we note that non-equilibrium models of simple repression have been shown to have the same functional form that we derive below [31]). We extend these models to consider the role of allostery by accounting for the equilibrium state of the repressor through the MWC model as follows.

Consider a cell with copy number *P* of RNAP and *R* repressors. Our model assumes that the repressor can exist in two conformational states. *R_A_* repressors will be in the active state (the favored state when the repressor is not bound to an inducer; in this state the repressor binds tightly to the DNA) and the remaining *R_I_* repressors will be in the inactive state (the predominant state when repressor is bound to an inducer; in this state the repressor binds weakly to the DNA) such that *R_A_* + *R_I_* = *R*. Repressors fluctuate between these two conformations in thermodynamic equilibrium [17].

Thermodynamic models of gene expression begin by enumerating all possible states of the promoter and their corresponding statistical weights. As shown in Fig. 2A, the promoter can either be empty, occupied by RNAP, or occupied by either an active or inactive repressor. We assign the repressor a different DNA binding affinity in the active and inactive state. In addition to the specific binding sites at the promoter, we assume that there are *N_NS_* non-specific binding sites elsewhere (i.e. on parts of the genome outside the simple repression architecture) where the RNAP or the repressor can bind. All specific binding energies are measured relative to the average non-specific binding energy. Our model explicitly ignores the complexity of the distribution of non-specific binding affinities in the genome, and makes the assumption that a single parameter can capture the energy difference between our binding site of interest and the average site in the reservoir. Thus, Δ*ε_P_* represents the energy difference between the specific and non-specific binding for RNAP to the DNA. Likewise, Δ*ε_RA_* and Δ*ε_RI_* represent the difference in specific and non-specific binding energies for repressor in the active or inactive state, respectively.

Thermodynamic models of transcription [2, 11, 14–16, 18, 27–30] posit that gene expression is proportional to the probability that the RNAP is bound to the promoter *p*_bound_, which is given by

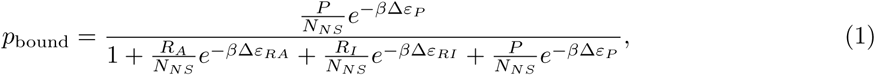

with 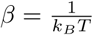 where *k_B_* is the Boltzmann constant and *T* is the temperature of the system. As *k_B_T* is the natural unit of energy at the molecular length scale, we treat the products *β*Δ*ε_j_* as single parameters within our model. Measuring *p*_bound_ directly is fraught with experimental difficulties, as determining the exact proportionality between expression and *p*_bound_ is not straightforward. Instead, we measure the fold-change in gene expression due to the presence of the repressor. We define fold-change as the ratio of gene expression in the presence of repressor relative to expression in the absence of repressor (i.e. constitutive expression), namely,

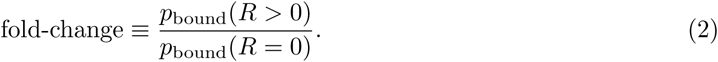

We can simplify this expression using two well-justified approximations: (1)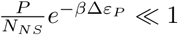 implying that the RNAP binds weakly to the promoter (*N_NS_* = 4.6×10^6^, *P* ≈ 10^3^ [32], Δ*ε_P_* ≈ −2 to −5 *k_B_T* [20], so that 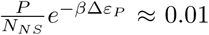) and (2) 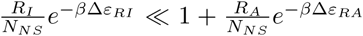 which reflects our assumption that the inactive repressor binds weakly to the promoter of interest. Using these approximations, the fold-change reduces to the form

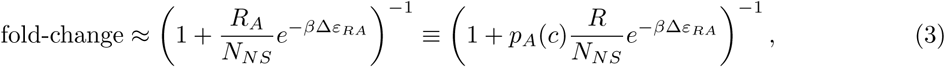

where in the last step we have introduced the fraction *p_A_*(*c*) of repressors in the active state given a concentration *c* of inducer, which is defined as *R_A_*(*c*) = *p_A_*(*c*)*R*. Since inducer binding shifts the repressors from the active to the inactive state, *p_A_*(*c*) is a decreasing function of *c* [10].

We compute the probability *p_A_*(*c*) that a repressor with *n* inducer binding sites will be active using the MWC model. After first enumerating all possible configurations of a repressor bound to inducer (see Fig. 2B), *p_A_*(*c*) is given by the sum of the weights of the active states divided by the sum of the weights of all possible states, namely,

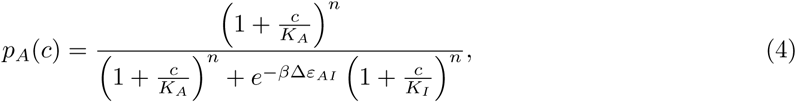

where *K_A_* and *K_I_* represent the dissociation constant between the inducer and repressor in the active and inactive states, respectively, and Δ*ε_AI_* = *ε_I_* − *ε_A_* stands for the free energy difference between a repressor in the inactive and active state (the quantity 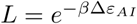 is sometimes denoted by *L* [10, 17] or *K*_RR*_ [11]). A repressor which favors the active state in the absence of inducer (Δ*ε_AI_* > 0) will be driven towards the inactive state upon inducer binding when *K_I_* < *K_A_*. The specific case of a repressor dimer with *n* = 2 inducer binding sites is shown in Fig. 2B.

Substituting *p_A_*(*c*) from Eq. (4) into Eq. (3) yields the general formula for induction of a simple repression regulatory architecture, namely,

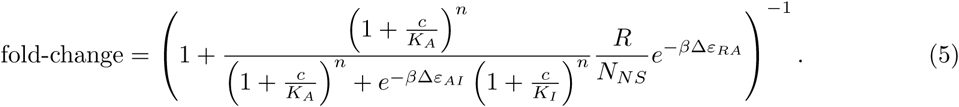

While we have used the specific case of simple repression with induction to craft this model, we reiterate that the exact same mathematics describe the case of corepression in which binding of an allosteric effector stabilizes the active state of the repressor and decreases gene expression (see Fig. 1B). Interestingly, we shift from induction (governed by *K_I_* < *K_A_*) to corepression (*K_I_* > *K_A_*) as the ligand transitions from preferentially binding to the inactive repressor state to stabilizing the active state. Furthermore, this general approach can be used to describe a variety of other motifs such as activation, multiple repressor binding sites, and combinations of activator and repressor binding sites [15, 16, 18].

This key formula presented in Eq. (5) enables us to make precise quantitative statements about induction profiles. Motivated by the broad range of predictions implied by this equation, we designed a series of experiments using the *lac* system in *E. coli* to tune the control parameters for a simple repression genetic circuit. As discussed in Fig. 3, previous studies from our lab have provided us with well-characterized values for many of the parameters in our experimental system, leaving only the values of the the MWC parameters (*K_A_*, *K_I_*, and Δ*ε_AI_*) to be determined. We note that while previous studies have obtained values for *K_A_*, *K_I_*, and 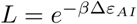 [11, 33], they were either based upon clever biochemical experiments or *in vivo* conditions involving poorly characterized transcription factor copy numbers and gene copy numbers. These differences relative to our experimental conditions and fitting techniques led us to believe that it was important to perform our own analysis of these parameters. Indeed, after inferring these three MWC parameters (see Appendix A for details regarding the inference of Δ*ε_AI_*, which was fitted separately from *K_A_* and *K_I_*), we were able to predict the input/output response of the system under a broad range of experimental conditions. For example, this framework can predict the response of the system at different repressor copy numbers *R*, repressor-operator affinities Δ*ε_RA_*, inducer concentrations *c*, and gene copy numbers (see Appendix B).

## Experimental Design

To test this model of allostery, we build off of a collection of work that has developed both a quantitative understanding of and experimental control over the simple repression motif. As shown in Fig. 3, earlier work from our laboratory used *E. coli* constructs based on components of the *lac* system to demonstrate how the Lac repressor (LacI) copy number *R* and operator binding energy Δ*ε_RA_* affect gene expression in the absence of inducer [14]. Rydenfelt *et al.* [34] extended the theory used in that work to the case of multiple promoters competing for a given transcription factor, which was demonstrated experimentally by Brewster *et al.* [15], who modified this system to consider expression from multiple-copy plasmids as well as the presence of competing repressor binding sites. Although the current work focuses on systems with a single site of repression, in Appendix A we utilize data from Brewster *et al.* [15] to characterize the allosteric free energy difference Δ*ε_AI_* between the repressor’s active and inactive states. With this parameter in hand, the present work considers the effects of an inducer on gene expression, adding yet another means for tuning the behavior of the system. A remarkable feature of our approach is how accurately our simple model quantitatively describes the mean response of a wide variety of regulatory contexts. We extend this body of work by introducing three additional biophysical parameters – Δ*ε_AI_*, *K_A_*, and *K_I_* – which capture the allosteric nature of the transcription factor and complement the results shown by Garcia and Phillips [14] and Brewster *et al.* [15].

To test this extension to the theory of transcriptional regulation by simple repression, we predicted the induction profiles for an array of strains that could be made using the previously characterized repressor copy number and DNA binding energies. More specifically, we used modified *lacI* ribosomal binding sites from Garcia and Phillips [14] to generate strains with mean repressor copy number per cell of *R* = 22 ± 4, 60 ± 20, 124 ± 30, 260 ± 40, 1220 ± 160, and 1740 ± 340, where the error denotes standard deviation of at least three replicates as measured by Garcia and Phillips [14]. We note that repressor copy number *R* refers to the number of repressor dimers in the cell, which is twice the number of repressor tetramers reported by Garcia and Phillips [14]; since both heads of the repressor are assumed to always be either specifically or non-specifically bound to the genome, the two repressor dimers in each LacI tetramer can be considered independently. Gene expression was measured using a Yellow Fluorescent Protein (YFP) gene, driven by a *lacUV5* promoter. Each of the six repressor copy number variants were paired with the native O1, O2, or O3 LacI operator [35] placed at the YFP transcription start site, thereby generating eighteen unique strains. The repressor-operator binding energies (O1 Δ*ε_RA_* = −15.3 ± 0.2 *k_B_T*, O2 Δ*ε_RA_* = −13.9 *k_B_T* ± 0.2, and O3 Δ*ε_RA_* = −9.7 ± 0.1 *k_B_ T*) were previously inferred by measuring the fold-change of the *lac* system at different repressor copy numbers, where the error arises from model fitting [14]. Additionally, we were able to obtain the value Δ*ε_AI_* = 4.5 *k_B_T* by fitting to previous data as discussed in Appendix A. We measure fold-change over a range of known IPTG concentrations *c*, using *n* = 2 inducer binding sites per LacI dimer and approximating the number of non-specific binding sites as the length in base-pairs of the *E. coli* genome, *N_NS_* = 4.6 × 10^6^. We proceed by first inferring the values of the repressor-inducer dissociation constants *K_A_* and *K_I_* using Bayesian inferential methods as discussed below [36]. When combined with the previously measured parameters within Eq. (5), this enables us to predict gene expression for any concentration of inducer, repressor copy number, and DNA binding energy.

Our experimental pipeline for determining fold-change using flow cytometry is shown in Fig. 4. Briefly, cells were grown to exponential phase, in which gene expression reaches steady state [37], under concentrations of the inducer IPTG ranging between 0 and 5 mM. We measure YFP fluorescence using flow cytometry and automatically gate the data to include only single-cell measurements (see Appendix C). To validate the use of flow cytometry, we also measured the fold-change of a subset of strains using the established method of single-cell microscopy (see Appendix D). We found that the fold-change measurements obtained from microscopy were indistinguishable from that of flow-cytometry and yielded values for the inducer binding constants *K_A_* and *K_I_* that were within error.

**Figure 4.**
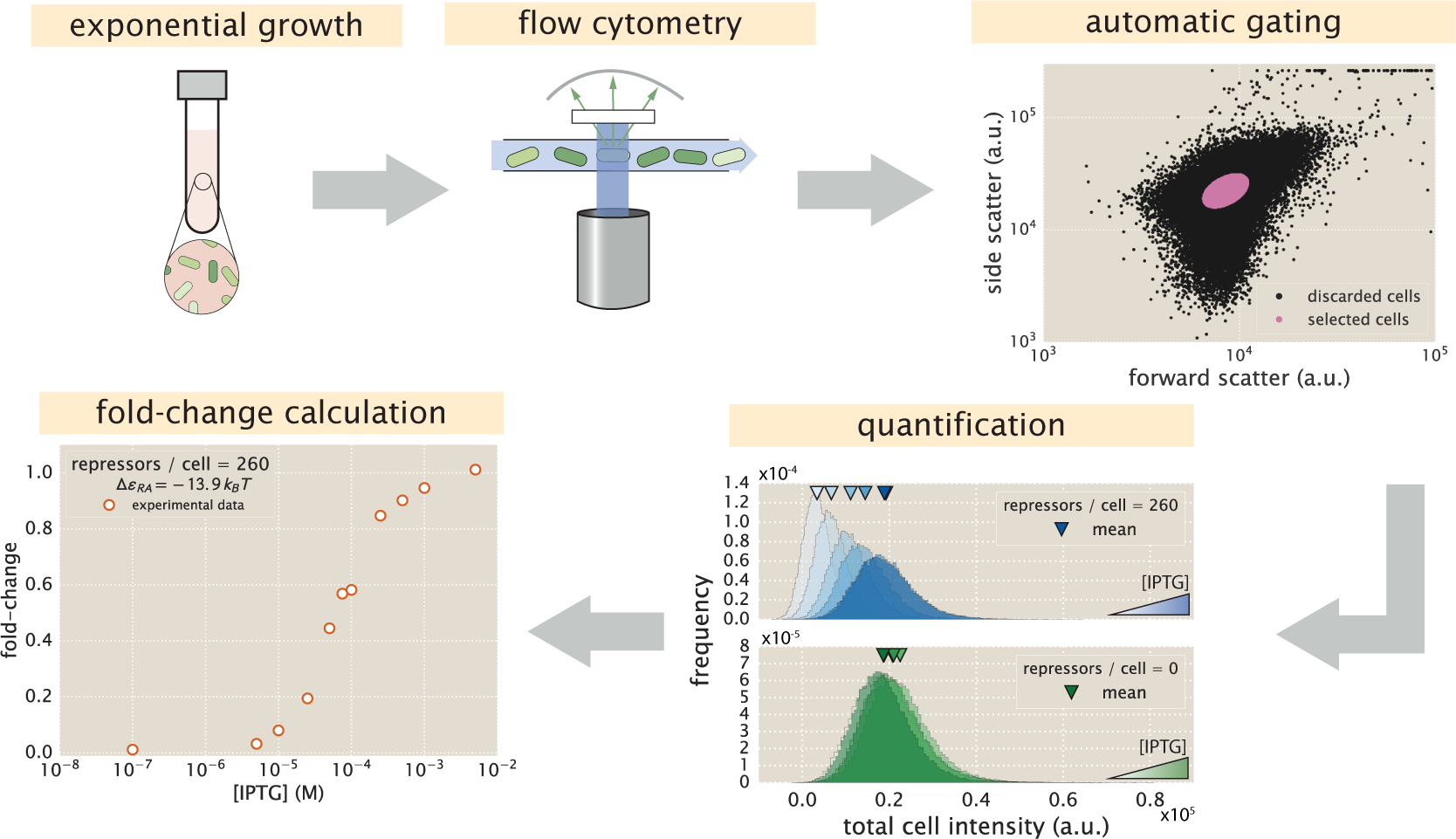
An experimental pipeline for high-throughput fold-change measurements. Cells are grown to exponential steady state and their fluorescence is measured using flow cytometry. Automatic gating methods using forward- and side-scattering are used to ensure that all measurements come from single cells (see Methods). Mean expression is then quantified at different IPTG concentrations (top, blue histograms) and for a strain without repressor (bottom, green histograms), which shows no response to IPTG as expected. Fold-change is computed by dividing the mean fluorescence in the presence of repressor by the mean fluorescence in the absence of repressor.

## Determination of the *in vivo* MWC Parameters

The three parameters that we tune experimentally are shown in Fig. 5A, leaving the three allosteric parameters (Δ*ε_AI_*, *K_A_*, and *K_I_*) to be determined by fitting. Using previous LacI fold-change data [15], we infer that Δ*ε_AI_* = 4.5 *k_B_T* (see Appendix A). Rather than fitting *K_A_* and *K_I_* to our entire data set of eighteen unique constructs, we performed a Bayesian parameter estimation on the data from a single strain with *R* = 260 and an O2 operator (Δ*ε_RA_* = −13.9 *k_B_T* [14]) shown in Fig. 5D (white circles). Using Markov Chain Monte Carlo, we determine the most likely parameter values to be 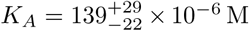 and 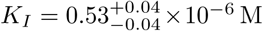, which are the modes of their respective distributions, where the superscripts and subscripts represent the upper and lower bounds of the 95^th^ percentile of the parameter value distributions as depicted in Fig. 5B. Unfortunately, we are not able to make a meaningful value-for-value comparison of our parameters to those of earlier studies [3, 11] because of the effects induced by uncertainties in both gene copy number and transcription factor numbers, the importance of which is illustrated by the plots in Appendix B. To demonstrate the strength of our parameter-free model, we then predicted the fold-change for the remaining seventeen strains with no further fitting (see Fig. 5C-E) together with the specific phenotypic properties described in Fig. 1 and discussed in detail below (see Fig. 5F-J). The shaded regions in Fig. 5C-J denote the 95% credible regions. An interesting aspect of our predictions of fold-change is that the width of the credible regions increases with repressor copy number and inducer concentration but decreases with the repressor-operator binding strength. Note that the fold-change Eq. (5) depends on the product of 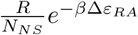 with the MWC parameters *K^A^*, *K^I^*, and Δ*ε_AI_*. As a result, strains with small repressor copy numbers, as well as strains with weak binding energies such as O3, will necessarily suppress variation in the MWC parameters (see Appendix E).

**Figure 5.**
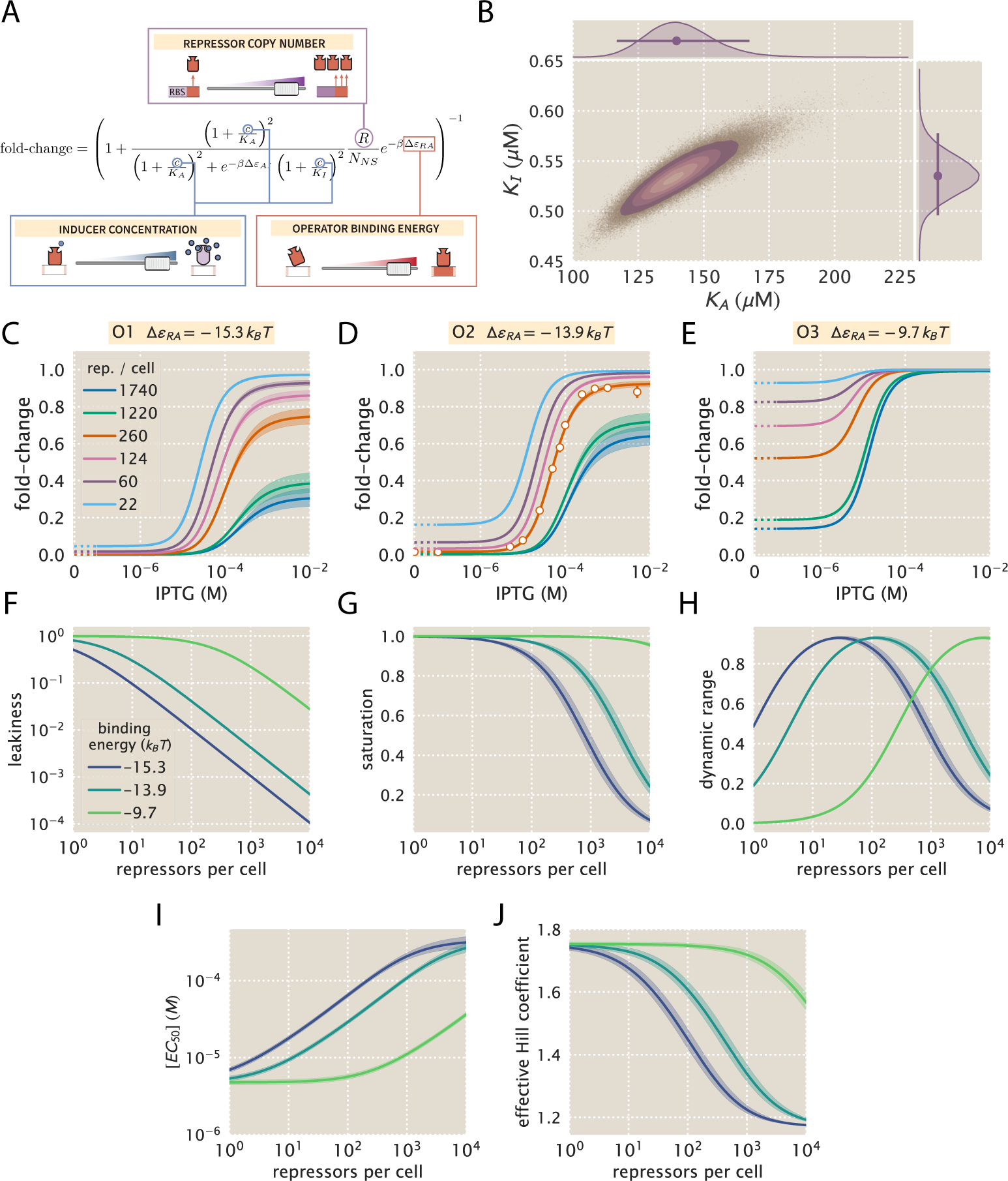
Predicting induction profiles for different biological control parameters. (A) We can quantitatively tune *R* via ribosomal binding site (RBS) modifications, Δ*ε_RA_* by mutating the operator sequence, and *c* by adding different amounts of IPTG to the growth medium. (B) Previous experiments have characterized the *R*, *N_NS_*, Δ*ε_RA_*, and Δ*ε_AI_* parameters (see Fig. 3), leaving only the unknown dissociation constants *K_A_* and *K_I_* between the inducer and the repressor in the active and inactive states, respectively. These two parameters can be inferred using Bayesian parameter estimation from a single induction curve. (C-E) Predicted IPTG titration curves for different repressor copy numbers and operator strengths. Titration data for the O2 strain (white circles in Panel D) with *R* = 260, Δ*ε_RA_* = −13.9 *k_B_T*, *n* = 2, and Δ*ε_AI_* = 4.5 *k_B_T* can be used to determine the thermodynamic parameters 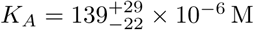 and 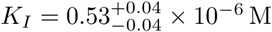 (orange line). The remaining solid lines predict the fold-change Eq. (5) for all other combinations of repressor copy numbers (shown in the legend) and repressor-DNA binding energies corresponding to the O1 operator (−15.3 *k_B_T*), O2 operator (−13.9 *k_B_T*), and O3 operator (−9.7 *k_B_T*). Error bars of experimental data show the standard error of the mean (eight or more replicates) when this error is not smaller than the diameter of the data point. The shaded regions denote the 95% credible region, although the credible region is obscured when it is thinner than the curve itself. To display the measured fold-change in the absence of inducer, we alter the scaling of the *x*-axis between 0 and 10^−7^ M to linear rather than logarithmic, as indicated by a dashed line. Additionally, our model allows us to investigate key phenotypic properties of the induction profiles (see Fig. 1B). Specifically, we show predictions for the (F) leakiness, (G) saturation, (H) dynamic range, (I) [*EC*_50_], and (J) effective Hill coefficient of the induction profiles.

We stress that the entire suite of predictions in Fig. 5 is based upon the induction profile of a single strain. Our ability to make such a broad range of predictions stems from the fact that our parameters of interest - such as the repressor copy number and DNA binding energy - appear as distinct physical parameters within our model. While the single data set in Fig. 5D could also be fit using a Hill function, such an analysis would be unable to predict any of the other curves in the figure. Phenomenological expressions such as the Hill function can describe data, but lack predictive power and are thus unable to build our intuition, design *de novo* input-output functions, or guide future experiments.

## Comparison of Experimental Measurements with Theoretical Predictions

We tested the predictions shown in Fig. 5 by measuring the fold-change induction profiles using strains that span this broad range in repressor copy numbers and repressor binding energies as characterized in [14], and inducer concentrations spanning several orders of magnitude. The results, shown in Fig. 6, demonstrate very good agreement between theory and experiment across all of our strains. We note, however, that there was an apparently systematic shift in the O3 Δ*ε_RA_* = −9.7 *k_B_T* strains (Fig. 6C) and all of the *R* = 1220 and *R* = 1740 strains. This may be partially due to imprecise previous determinations of their Δ*ε_RA_* and *R* values. By performing a global fit where we infer all parameters including the repressor copy number *R* and the binding energy Δ*ε_RA_*, we found better agreement for these particular strains, although a discrepancy in the steepness of the response for all O3 strains remains (see Appendix F). As an additional test of our model, we also considered strains using the synthetic Oid operator which exhibits stronger repression, Δ*ε_RA_* = −17 *k_B_T* [14], than the O1, O2, and O3 operators. We found that we were unable to measure the strongly repressed strains accurately by flow cytometry. However, for the data we collected, we found that the MWC description was consistent to within 1 *k_B_T* of the binding energy previously reported (see Appendix G for more details).

**Figure 6.**
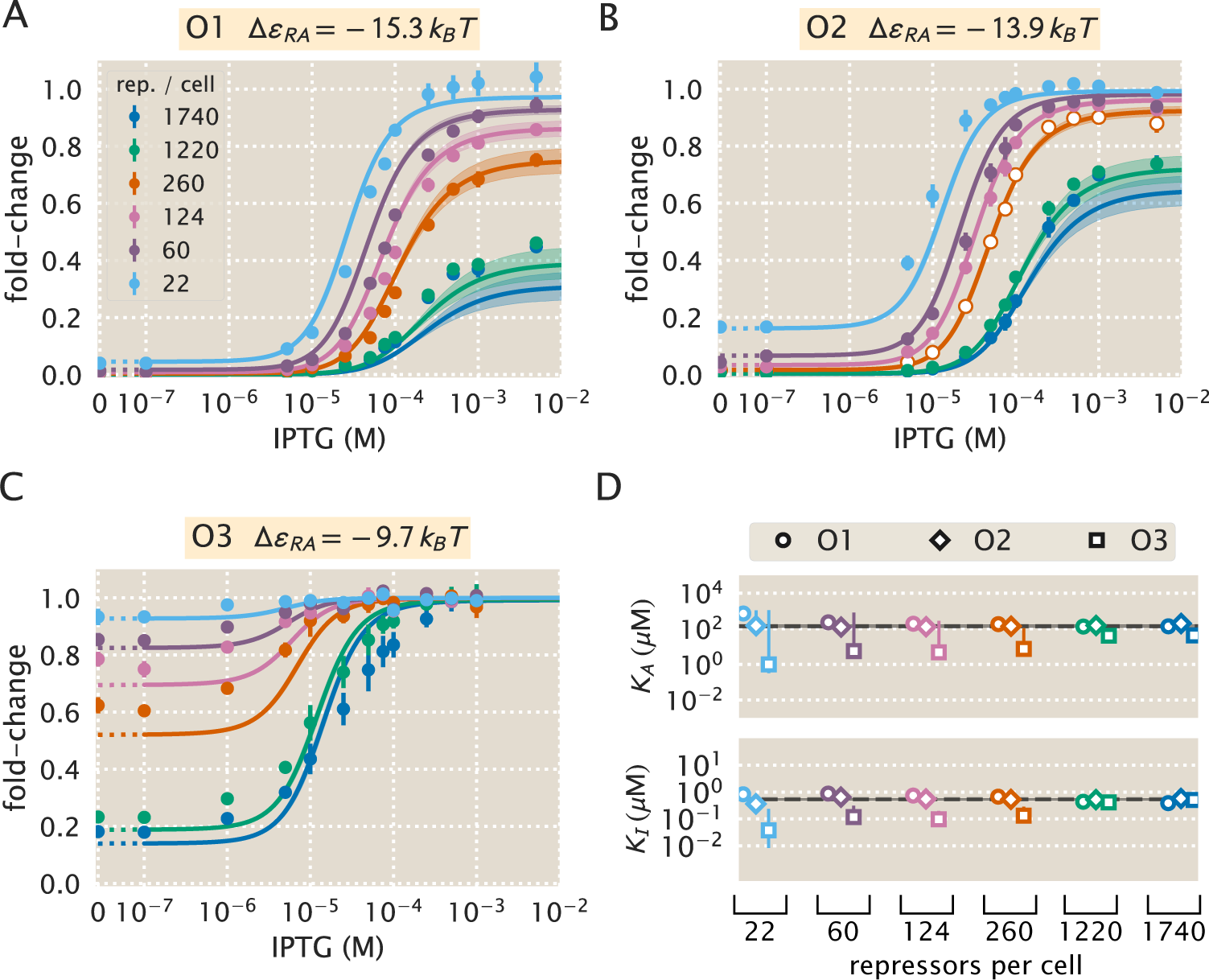
Comparison of predictions against measured and inferred data. Flow cytometry measurements of fold-change over a range of IPTG concentrations for (A) O1, (B) O2, and (C) O3 strains at varying repressor copy numbers, overlaid on the predicted responses. Error bars of the experimental data show the standard error of the mean (eight or more replicates). As discussed in Fig. 5, all of the predicted induction curves were created prior to measurement by inferring the MWC parameters using a single data set (O2 *R* = 260, shown by white circles in Panel B). The predictions may therefore depend upon which strain is used to infer the parameters. (D) The inferred parameter values of the dissociation constants *K_A_* and *K_I_* using any of the eighteen strains instead of the O2 *R* = 260 strain. Nearly identical parameter values are inferred from each strain, demonstrating that the same set of induction profiles would have been predicted regardless of which strain was chosen. The points show the mode and the error bars denote the 95% credible region of the parameter value distribution. Error bars not visible are smaller than the size of the marker.

To ensure that the agreement between our predictions and data is not an accident of the strain we chose to perform our fitting, we explored the effects of using each of our other strains to estimate *K_A_* and *K_I_*. As shown in Appendix H and Fig. 6D, the inferred values of *K_A_* and *K_I_* depend very minimally upon which strain is chosen, demonstrating that these parameter values are highly robust. As previously mentioned, we performed a global fit using the data from all eighteen strains for the following parameters: the inducer dissociation constants *K_A_* and *K_I_*, the repressor copy numbers *R*, and the repressor DNA binding energy Δ*ε_RA_* (see Appendix F). This global fit led to very similar parameter values, lending strong support for our quantitative understanding of induction in the simple repression architecture. For the remainder of the text we proceed using our analysis on the strain with *R* = 260 repressors and an O2 operator.

## Predicting the Phenotypic Traits of the Induction Response

Rather than measuring the full induction response of a system, a subset of the properties shown in Fig. 1, namely, the leakiness, saturation, dynamic range, [*EC*_50_], and effective Hill coefficient, may be of greater interest. For example, synthetic biology is often focused on generating large responses (i.e. a large dynamic range) or finding a strong binding partner (i.e. a small [*EC*_50_]) [38, 39]. While these properties are all individually informative, when taken together they capture the essential features of the induction response. We reiterate that a Hill function approach cannot predict these features *a priori* and furthermore requires fitting each curve individually. The MWC model, on the other hand, enables us to quantify how each trait depends upon a single set of physical parameters as shown by Fig. 5F-J.

We define these five phenotypic traits using expressions derived from the model, Eq. (5). These results build upon extensive work by Martins and Swain, who computed many such properties for ligand-receptor binding within the MWC model [9]. We begin by analyzing the leakiness, which is the minimum fold-change observed in the absence of ligand, given by

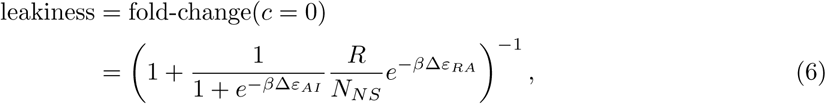

and the saturation, which is the maximum fold change observed in the presence of saturating ligand,

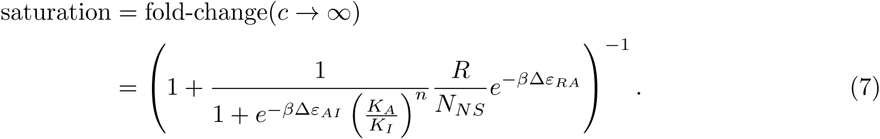

Systems that minimize leakiness repress strongly in the absence of effector while systems that maximize saturation have high expression in the presence of effector. Together, these two properties determine the dynamic range of a system’s response, which is given by the difference

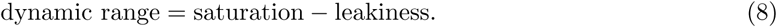

These three properties are shown in Fig. 5F-H. We discuss these properties in greater detail in Appendix I. For example, we compute the number of repressors *R* necessary to evoke the maximum dynamic range and demonstrate that the magnitude of this maximum is independent of the repressor-operator binding energy Δ*ε_RA_*. Fig. 7A-C show that the measurements of these three properties, derived from the fold-change data in the absence of IPTG and the presence of saturating IPTG, closely match the predictions for all three operators.

**Figure 7.**
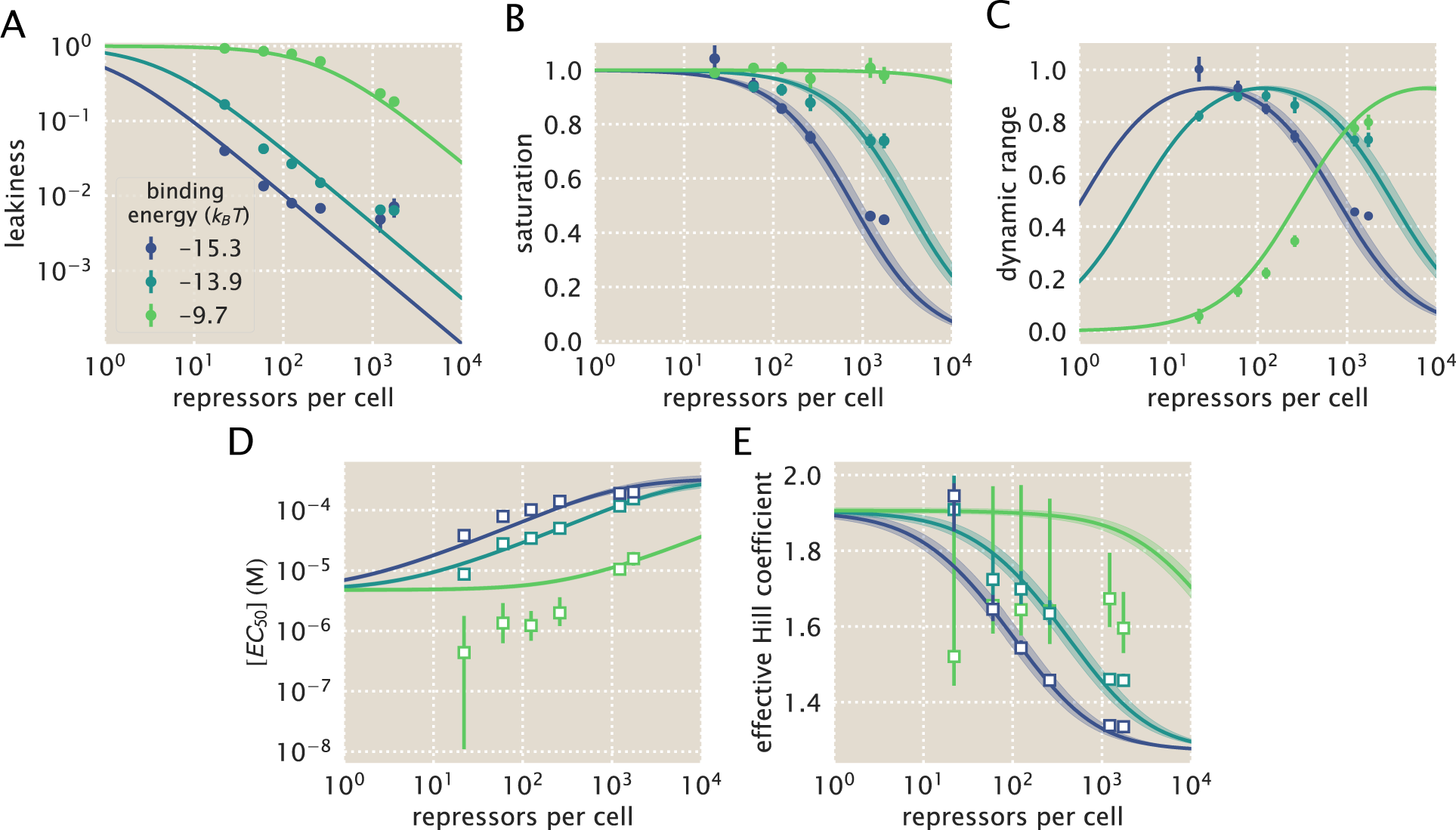
Predictions and experimental measurements of key properties of induction profiles. Data for the (A) leakiness, (B) saturation, and (C) dynamic range are obtained from fold-change measurements in Fig. 6 in the absence of IPTG and at saturating concentrations of IPTG. The three repressor-operator binding energies in the legend correspond to the O1 operator (−15.3 *k_B_T*), O2 operator (−13.9 *k_B_T*), and O3 operator (−9.7 *k_B_T*). Both the (D) [*EC*_50_] and (E) effective Hill coefficient are inferred by individually fitting each operator-repressor pairing in Fig. 6A-C separately to Eq. (5) in order to smoothly interpolate between the data points. Error bars for A-C represent the standard error of the mean for eight or more replicates; error bars for D-E represent the 95% credible region for the parameter found by propagating the credible region of our estimates of *K_A_* and *K_I_* into Eqs. (9) and (10).

Two additional properties of induction profiles are the [*EC*_50_] and effective Hill coefficient, which determine the range of inducer concentration in which the system’s output goes from its minimum to maximum value. The [*EC*_50_] denotes the inducer concentration required to generate a system response Eq. (5) halfway between its minimum and maximum value,

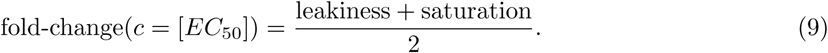

The effective Hill coefficient *h*, which quantifies the steepness of the curve at the [*EC*_50_] [10], is given by

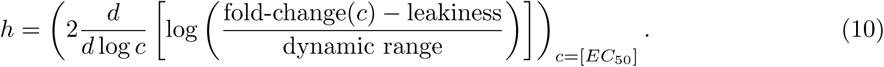

Fig. 5I-J shows how the [*EC*_50_] and effective Hill coefficient depend on the repressor copy number. In Appendix I, we discuss the analytic forms of these two properties as well as their dependence on the repressor-DNA binding energy.

Fig. 7D-E show the estimated values of the [*EC*_50_] and the effective Hill coefficient overlaid on the theoretical predictions. Both properties were obtained by fitting Eq. (5) to each individual titration curve and computing the [*EC*_50_] and effective Hill coefficient using Eq. (9) and Eq. (10), respectively. We find that the predictions made with the single strain fit closely match those made for each of the strains with O1 and O2 operators, but the predictions for the O3 operator are markedly off. The large, asymmetric error bars for the O3 *R* = 22 strain arise from its nearly flat response, where the lack of dynamic range makes it impossible to determine the value of the inducer dissociation constants *K_A_* and *K_I_*; consequently the determination of [*EC*_50_] is accompanied with significant uncertainty.

## Data Collapse of Induction Profiles

Our primary interest heretofore was to determine the system response at a specific inducer concentration, repressor copy number, and repressor-DNA binding energy. We now flip this question on its head and ask: given a specific value of the fold-change, what combination of parameters will give rise to this desired response? In other words, what trade-offs between the parameters of the system will give rise to the same mean cellular output? These are key questions both for understanding how the system is governed and for engineering specific responses in a synthetic biology context. To this end, we rewrite Eq. (5) as a Fermi function,

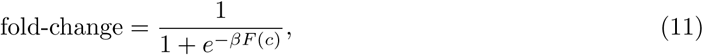

where *F*(*c*) is the free energy of the repressor binding to the operator of interest relative to the unbound operator state [23, 24, 31], which is given by

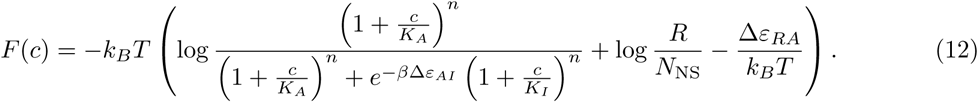

The first term in the parenthesis denotes the contribution from the inducer concentration, the second the effect of the repressor copy number, and the last the repressor-operator binding energy. We note that elsewhere, this free energy has been dubbed the Bohr parameter since such families of curves are analogous to the shifts in hemoglobin binding curves at different pHs known as the Bohr effect [31, 40, 41].

Instead of analyzing each induction curve individually, the free energy provides a natural means to simultaneously characterize the diversity in our eighteen induction profiles. Fig. 8A demonstrates how the various induction curves from Fig. 5C-E all collapse onto a single master curve, where points from every induction profile that yield the same fold-change are mapped onto the same free energy. Fig. 8B shows this data collapse for the 216 data points in Fig. 6A-C, demonstrating the close match between the theoretical predictions and experimental measurements across all eighteen strains.

**Figure 8.**
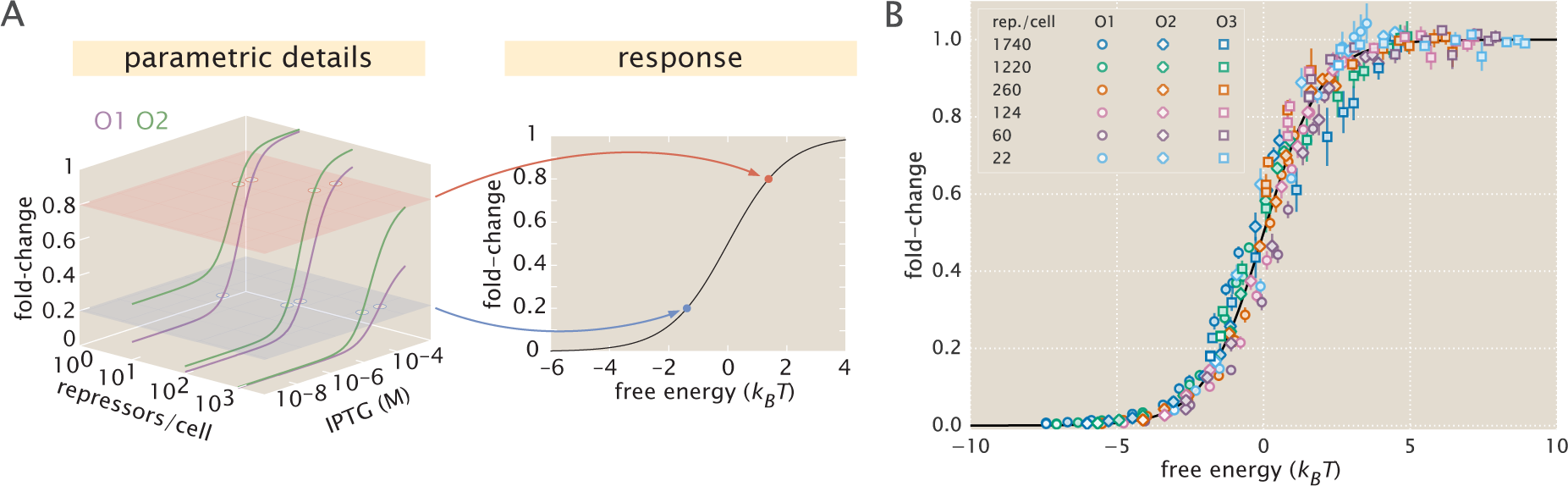
Fold-change data from a broad collection of different strains collapse onto a single master curve. (A) Any combination of parameters can be mapped to a single physiological response (i.e. fold-change) via the free energy, which encompasses the parametric details of the model. (B) Experimental data from Fig. 6 collapse onto a single master curve as a function of the free energy Eq. (12). The free energy for each strain was calculated from Eq. (12) using *n* = 2, Δ*ε_AI_* = 4.5 *k_B_T*, *K_A_* = 139 × 10^−6^ M, *K_I_* = 0.53 × 10^−6^ M, and the strain specific *R* and Δ*ε_RA_*. All data points represent the mean and error bars are the standard error of the mean for eight or more replicates.

There are many different combinations of parameter values that can result in the same free energy as defined in Eq. (12). For example, suppose a system originally has a fold-change of 0.2 at a specific inducer concentration, and then operator mutations increase the Δ*ε_RA_* binding energy. While this serves to initially increase both the free energy and the fold-change, a subsequent increase in the repressor copy number could bring the cell back to the original fold-change level. Such trade-offs hint that there need not be a single set of parameters that evoke a specific cellular response, but rather that the cell explores a large but degenerate space of parameters with multiple, equally valid paths.

## Discussion

Since the early work by Monod, Wyman, and Changeux [17, 42], a broad list of different biological phenomena have been tied to the existence of macromolecules that switch between inactive and active states. Examples can be found in a wide variety of cellular processes that include ligand-gated ion channels [43], enzymatic reactions [41, 44], chemotaxis [24], quorum sensing [23], G-protein coupled receptors [45], physiologically important proteins [46, 47], and beyond. One of the most ubiquitous examples of allostery is in the context of gene expression, where an array of molecular players bind to transcription factors to either aid or deter their ability to regulate gene activity [25, 26]. Nevertheless, no definitive study has been made of the applicability of the MWC model to transcription factor function, despite the clear presence of different conformational states in their structures in the presence and absence of signaling molecules [48]. A central goal of this work was to assess whether a thermodynamic MWC model can provide an accurate input-output function for gene regulation by allosteric transcription factors.

Others have developed quantitative models describing different aspects of allosteric regulatory systems. Martins and Swain analytically derived fundamental properties of the MWC model, including the leakiness and dynamic range described in this work, noting the inherent trade-offs in these properties when tuning the microscopic parameters of the model [9, 10]. Work in the Church and Voigt labs, among others, has expanded on the availability of allosteric circuits for synthetic biology [7, 8, 49, 50]. Recently, Daber *et al.* theoretically explored the induction of simple repression within the MWC model [3] and experimentally measured how mutations alter the induction profiles of transcription factors [11]. Vilar and Saiz considered the broad range of interactions in inducible *lac*-based systems including the effects of oligomerization and DNA folding on transcription factor induction [6, 51]. Other work has attempted to use the *lac* system to reconcile *in vitro* and *in vivo* measurements [12, 52]. Although this body of work has done much to improve our understanding of allosteric transcription factors, there has remained a disconnect between model and experiment. In order to rigorously test a model’s applicability to natural systems, the model’s predictions must be weighed against data from precise experiments specifically designed to test those predictions.

Here, we expand upon this body of work by generating a predictive model of allosteric transcriptional regulation and then testing the model against a thorough set of experiments using well-characterized regulatory components. Specifically, we used the MWC model to build upon and refine a well-established thermodynamic model of transcriptional regulation [14, 18], allowing us to compose the model from a minimal set of biologically meaningful parameters. This minimal model captures the key players of transcriptional regulation – namely the repressor copy number, the DNA binding energy, and the concentration of inducer – and enables us to predict how the system will behave when we change each of these parameters. We tested these predictions on a range of strains whose repressor copy number spanned two orders of magnitude and whose DNA binding affinity spanned 6 *k_B_T*. We argue that one would not be able to generate such a wide array of predictions by using a Hill function, which abstracts away the biophysical meaning of the parameters into phenomenological parameters [53].

Specifically, we tested our model in the context of a *lac*-based simple repression system by first determining the allosteric dissociation constants *K_A_* and *K_I_* from a single induction data set (O2 operator with binding energy Δ*ε_RA_* = −13.9 *k_B_T* and repressor copy number *R* = 260) and then using these values to make parameter-free predictions of the induction profiles for seventeen other strains where Δ*ε_RA_* and *R* were varied significantly (see Fig. 5). We next measured the induction profiles of these seventeen strains using flow cytometry and found that our predictions consistently and accurately captured the primary features for each induction data set, as shown in Fig. 6A-C. Surprisingly, we find that the inferences for the repressor-inducer dissociation constants that would have been derived from any other single strain (instead of the O2 operator with *R* = 260) would have resulted in nearly identical predictions (see Fig. 6D and Appendix H). This suggests that a few carefully chosen measurements can lead to a deep quantitative understanding of how simple regulatory systems work without requiring an extensive sampling of strains that span the parameter space. Moreover, the fact that we could consistently achieve reliable predictions after fitting only two free parameters stands in contrast to the common practice of fitting several free parameters simultaneously, which can nearly guarantee an acceptable fit provided that the model roughly resembles the system response, regardless of whether the details of the model are tied to any underlying molecular mechanism.

Beyond observing changes in fold-change as a function of effector concentration, our application of the MWC model allows us to explicitly predict the values of the induction curves’ key parameters, namely, the leakiness, saturation, dynamic range, [*EC*_50_], and the effective Hill coefficient (see Fig. 7). This allows us to quantify the unique traits of each set of strains examined here. Strains using the O1 operator consistently have a low leakiness value, a consequence of its strong binding energy. The saturation values for these strains, however, vary significantly with *R*. This trend is reversed for strains using O3, which has the weakest binding energy of our constructs. Leakiness values for constructs using O3 vary strongly with *R*, but their saturation values approach 1 regardless of *R*. Strains with the intermediate O2 binding energy have both a leakiness and saturation that vary markedly with *R*. For both the O1 and O2 data sets, our model also accurately predicts the effective Hill coefficient and [*EC*_50_], though these predictions for O3 are noticeably less accurate. While performing a global fit for all model parameters marginally improves the prediction for O3 (see Appendix F), we are still unable to accurately predict the effective Hill coefficient or the [*EC*_50_], though the uncertainties in these two parameters are really an inheritance from the consistent difference between the theoretical and measured sharpness of the induction response seen in Fig. 6C.

Because this model allows us to derive expressions for individual features of induction curves, we are able to examine how these features may be tuned by careful selection of system parameters. Fig. 7 shows how each of the induction curves’ key features vary as a function of Δ*ε_RA_* and *R*, which makes it possible to select desired properties from among the possible phenotypes available to the system. For instance, it is possible to obtain a high dynamic range using fewer than 100 repressors if the binding energy is strong. As an example of the constraints inherent to the system, one cannot design a strain with a leakiness of 0.1 and a saturation of 0.4 by only varying the repressor copy number and repressor-operator binding affinity, since these two properties are coupled by Eqs. (6) and (7). Achieving this particular behavior would require changing the ratio *K_A_/K_I_* of repressor-inducer dissociation constants, as may be done by mutating the repressor’s inducer binding pocket.

The dynamic range, which is of considerable interest when designing or characterizing a genetic circuit, is revealed to have an interesting property: although changing the value of Δ*ε_RA_* causes the dynamic range curves to shift to the right or left, each curve has the same shape and in particular the same maximum value. This means that strains with strong or weak binding energies can attain the same dynamic range when the value of *R* is tuned to compensate for this energy. This feature is not immediately apparent from the IPTG induction curves, which show very low dynamic ranges for several of the O1 and O3 strains. Without the benefit of models that can predict such phenotypic traits, efforts to engineer genetic circuits with allosteric transcription factors must rely on trial and error to achieve specific responses [7, 8]. This is a compelling example showing that our predictive modeling approach has a significant advantage over descriptive models.

To our knowledge this is the first work of its kind in which a single family of parameters is demonstrated to predict a vast range of induction curves with qualitatively different behaviors. One of the demanding criteria of our approach is that a small set of parameters must consistently describe data from a diverse collection of data sets taken using distinct methods such as Miller assays and bulk and single-cell fluorescence experiments to measure fold-change (see Appendices C and G), as well as quantitative Western blots [14] and binomial partitioning methods to count repressors [15, 54]. Furthermore, we build off of our previous studies that use the simple repression architecture and we demand that the parameters derived from these studies account for constructs that are integrated into the chromosome, plasmid-borne, and even for cases where there are competing binding sites to take repressors out of circulation [14, 15] (see Appendix B) or where there are multiple operators to allow DNA looping [21]. The resulting model not only predicts the individual titration profiles as a function of IPTG, but describes key properties of the response. The general agreement with the entire body of work presented here demonstrates that our model captures the underlying mechanism governing simple repression. We are unaware of any comparable study in transcriptional regulation that demands one predictive framework cover such a broad array of regulatory situations.

Despite the diversity observed in the induction profiles of each of our strains, our data are unified by their reliance on fundamental biophysical parameters. In particular, we have shown that our model for fold-change can be rewritten in terms of the free energy Eq. (12), which encompasses all of the physical parameters of the system. This has proven to be an illuminating technique in a number of studies of allosteric proteins [23, 24, 55]. Although it is experimentally straightforward to observe system responses to changes in effector concentration *c*, framing the input-output function in terms of *c* can give the misleading impression that changes in system parameters lead to fundamentally altered system responses. Alternatively, if one can find the “natural variable” that enables the output to collapse onto a single curve, it becomes clear that the system’s output is not governed by individual system parameters, but rather the contributions of multiple parameters that define the natural variable.

When our fold-change data are plotted against the respective free energies for each construct, they collapse cleanly onto a single curve (see Fig. 8). This enables us to analyze how parameters can compensate each other. For example, we may wish to determine which combinations of parameters result in a system that is strongly repressed (free energy *F*(*c*) ≤ −5 *k_B_T*). We know from our understanding of the induction phenomenon that strong repression is most likely to occur at low values of *c*. However, from Eq. (12) we can clearly see that increases in the value of *c* can be compensated by an increase in the number of repressors *R*, a decrease in the binding energy Δ*ε_RA_* (i.e. stronger binding), or some combination of both. Likewise, while the system tends to express strongly (*F*(*c*) ≥ 5 *k_B_T*) when *c* is high, one could design a system that expresses strongly at low values of *c* by reducing *R* or increasing the value of Δ*ε_RA_*. As a concrete example, given a concentration *c* = 10^−5^ M, a system using the O1 operator (Δ*ε_RA_* = −15.3 *k_B_T*) requires 745 or more repressors for *F*(*c*) ≤ −5 *k_B_ T*, while a system using the weaker O3 operator (Δ*ε_RA_* = −9.7 *k_B_ T*) requires 2 × 10^5^ or more repressors for *F* (*c*) ≤ −5 *k_B_T*.

While our experiments validated the theoretical predictions in the case of simple repression, we expect the framework presented here to apply much more generally to different biological instances of allosteric regulation. For example, we can use this model to explore different regulatory configurations such as corepression, activation, and coactivation, each of which are found in *E. coli* (see Appendix J). This work can also serve as a springboard to characterize not just the mean but the full gene expression distribution and thus quantify the impact of noise on this system [56]. Another extension of this approach would be to theoretically predict and experimentally verify whether the repressor-inducer dissociation constants *K_A_* and *K_I_* or the energy difference Δ*ε_AI_* between the allosteric states can be tuned by making single amino acid substitutions in the transcription factor [11, 31]. Finally, we expect that the kind of rigorous quantitative description of the allosteric phenomenon provided here will make it possible to construct biophysical models of fitness for allosteric proteins similar to those already invoked to explore the fitness effects of transcription factor binding site strengths and protein stability [57–59].

To conclude, we find that our application of the MWC model provides an accurate, predictive framework for understanding simple repression by allosteric transcription factors. To reach this conclusion, we analyzed the model in the context of a well-characterized system, in which each parameter had a clear biophysical meaning. As many of these parameters had been measured or inferred in previous studies, this gave us a minimal model with only two free parameters which we inferred from a single data set. We then accurately predicted the behavior of seventeen other data sets in which repressor copy number and repressor-DNA binding energy were systematically varied. In addition, our model allowed us to understand how key properties such as the leakiness, saturation, dynamic range, [*EC*_50_], and effective Hill coefficient depended upon the small set of parameters governing this system. Finally, we show that by framing inducible simple repression in terms of free energy, the data from all of our experimental strains collapse cleanly onto a single curve, illustrating the many ways in which a particular output can be targeted. In total, these results show that a thermodynamic formulation of the MWC model supersedes phenomenological fitting functions for understanding transcriptional regulation by allosteric proteins.

## Methods

### Bacterial Strains and DNA Constructs

All strains used in these experiments were derived from *E. coli* K12 MG1655 with the *lac* operon removed, adapted from those created and described in [14, 19]. Briefly, the operator variants and YFP reporter gene were cloned into a pZS25 background which contains a *lacUV5* promoter that drives expression as is shown in Fig. 2. These constructs carried a kanamycin resistance gene and were integrated into the *galK* locus of the chromosome using *λ* Red recombineering [60]. The *lacI* gene was constitutively expressed via a P_LtetO-1_ promoter [50], with ribosomal binding site mutations made to vary the LacI copy number as described in [61] using site-directed mutagenesis (Quickchange II; Stratagene), with further details in [14]. These *lacI* constructs carried a chloramphenicol resistance gene and were integrated into the *ybcN* locus of the chromosome. Final strain construction was achieved by performing repeated P1 transduction [62] of the different operator and *lacI* constructs to generate each combination used in this work. Integration was confirmed by PCR amplification of the replaced chromosomal region and by sequencing. Primers and final strain genotypes are listed in Appendix K.

It is important to note that the rest of the *lac* operon (*lacZYA*) was never expressed. The LacY protein is a transmembrane protein which actively transports lactose as well as IPTG into the cell. As LacY was never produced in our strains, we assume that the extracellular and intracellular IPTG concentration was approximately equal due to diffusion across the membrane into the cell as is suggested by previous work [63].

To make this theory applicable to transcription factors with any number of DNA binding domains, we used a different definition for repressor copy number than has been used previously. We define the LacI copy number as the average number of repressor dimers per cell whereas in [14], the copy number is defined as the average number of repressor tetramers in each cell. To motivate this decision, we consider the fact that the LacI repressor molecule exists as a tetramer in *E. coli* [48] in which a single DNA binding domain is formed from dimerization of LacI proteins, so that wild-type LacI might be described as dimer of dimers. Since each dimer is allosterically independent (i.e. either dimer can be allosterically active or inactive, independent of the configuration of the other dimer) [3], a single LacI tetramer can be treated as two functional repressors. Therefore, we have simply multiplied the number of repressors reported in [14] by a factor of two. This factor is included as a keyword argument in the numerous Python functions used to perform this analysis, as discussed in the code documentation.

A subset of strains in these experiments were measured using fluorescence microscopy for validation of the flow cytometry data and results. To aid in the high-fidelity segmentation of individual cells, the strains were modified to constitutively express an mCherry fluorophore. This reporter was cloned into a pZS4^*^1 backbone [50] in which mCherry is driven by the *lacUV5* promoter. All microscopy and flow cytometry experiments were performed using these strains.

### Growth Conditions for Flow Cytometry Measurements

All measurements were performed with *E. coli* cells grown to mid-exponential phase in standard M9 minimal media (M9 5X Salts, Sigma-Aldrich M6030; 2 mM magnesium sulfate, Mallinckrodt Chemicals 6066-04; 100 *μ*M calcium chloride, Fisher Chemicals C79-500) supplemented with 0.5% (w/v) glucose. Briefly, 500 *μ*L cultures of *E. coli* were inoculated into Lysogeny Broth (LB Miller Powder, BD Medical) from a 50% glycerol frozen stock (-80°C) and were grown overnight in a 2 mL 96-deep-well plate sealed with a breathable nylon cover (Lab Pak - Nitex Nylon, Sefar America Inc. Cat. No. 241205) with rapid agitation for proper aeration. After approximately 12 to 15 hours, the cultures had reached saturation and were diluted 1000-fold into a second 2 mL 96-deep-well plate where each well contained 500 *μ*L of M9 minimal media supplemented with 0.5% w/v glucose (anhydrous D-Glucose, Macron Chemicals) and the appropriate concentration of IPTG (Isopropyl *β*-D-1 thiogalactopyranoside Dioxane Free, Research Products International). These were sealed with a breathable cover and were allowed to grow for approximately eight hours. Cells were then diluted ten-fold into a round-bottom 96-well plate (Corning Cat. No. 3365) containing 90 *μ*L of M9 minimal media supplemented with 0.5% w/v glucose along with the corresponding IPTG concentrations. For each IPTG concentration, a stock of 100-fold concentrated IPTG in double distilled water was prepared and partitioned into 100 *μ*L aliquots. The same parent stock was used for all experiments described in this work.

### Flow Cytometry

Unless explicitly mentioned, all fold-change measurements were collected on a Miltenyi Biotec MACSquant Analyzer 10 Flow Cytometer graciously provided by the Pamela Björkman lab at Caltech. Detailed information regarding the voltage settings of the photo-multiplier detectors can be found in Appendix Table S1. Prior to each day’s experiments, the analyzer was calibrated using MACSQuant Calibration Beads (Cat. No. 130-093-607) such that day-to-day experiments would be comparable. All YFP fluorescence measurements were collected via 488 nm laser excitation coupled with a 525/50 nm emission filter. Unless otherwise specified, all measurements were taken over the course of two to three hours using automated sampling from a 96-well plate kept at approximately 4° - 10°C on a MACS Chill 96 Rack (Cat. No. 130-094-459). Cells were diluted to a final concentration of approximately 4 × 10^4^ cells per *μ*L which corresponded to a flow rate of 2,000-6,000 measurements per second, and acquisition for each well was halted after 100,000 events were detected. Once completed, the data were extracted and immediately processed using the following methods.

### Unsupervised Gating of Flow Cytometry Data

Flow cytometry data will frequently include a number of spurious events or other undesirable data points such as cell doublets and debris. The process of restricting the collected data set to those data determined to be “real” is commonly referred to as gating. These gates are typically drawn manually [64] and restrict the data set to those points which display a high degree of linear correlation between their forward-scatter (FSC) and side-scatter (SSC). The development of unbiased and unsupervised methods of drawing these gates is an active area of research [65, 66]. For our purposes, we assume that the fluorescence level of the population should be log-normally distributed about some mean value. With this assumption in place, we developed a method that allows us to restrict the data used to compute the mean fluorescence intensity of the population to the smallest two-dimensional region of the log(FSC) vs. log(SSC) space in which 40% of the data is found. This was performed by fitting a bivariate Gaussian distribution and restricting the data used for calculation to those that reside within the 40th percentile. This procedure is described in more detail in the supplementary information as well as in a Jupyter notebook located in this paper’s Github repository.

### Experimental Determination of Fold-Change

For each strain and IPTG concentration, the fold-change in gene expression was calculated by taking the ratio of the population mean YFP expression in the presence of LacI repressor to that of the population mean in the absence of LacI repressor. However, the measured fluorescence intensity of each cell also includes the autofluorescence contributed by the weak excitation of the myriad protein and small molecules within the cell. To correct for this background, we computed the fold change as

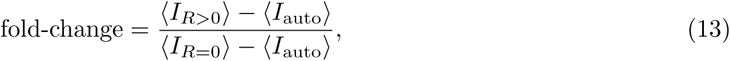

where 〈*I_R_*_>0_〉 is the average cell YFP intensity in the presence of repressor, 〈*I_R_*_=0_〉 is the average cell YFP intensity in the absence of repressor, and 〈*I*_auto_〉 is the average cell autofluorescence intensity, as measured from cells that lack the *lac*-YFP construct.

### Bayesian Parameter Estimation

In this work, we determine the the most-likely parameter values for the inducer dissociation constants *K_A_* and *K_I_* of the active and inactive state, respectively, using Bayesian methods. We compute the probability distribution of the value of each parameter given the data *D*, which by Bayes’ theorem is given by

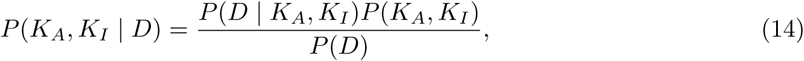

where *D* is all the data composed of independent variables (repressor copy number *R*, repressor-DNA binding energy Δ*ε_RA_*, and inducer concentration *c*) and one dependent variable (experimental fold-change). *P*(*D* | *K_A_, K_I_*) is the likelihood of having observed the data given the parameter values for the dissociation constants, *P*(*K_A_, K_I_*) contains all the prior information on these parameters, and *P* (*D*) serves as a normalization constant, which we can ignore in our parameter estimation. Eq. (5) assumes a deterministic relationship between the parameters and the data, so in order to construct a probabilistic relationship as required by Eq. (14), we assume that the experimental fold-change for the *i*^th^ datum given the parameters is of the form

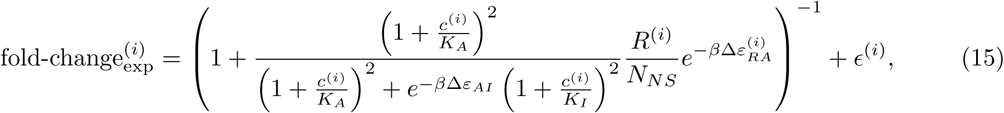

where *∊*^(*i*)^ represents the departure from the deterministic theoretical prediction for the *i*^th^ data point. If we assume that these *∊*^(*i*)^ errors are normally distributed with mean zero and standard deviation *σ*, the likelihood of the data given the parameters is of the form

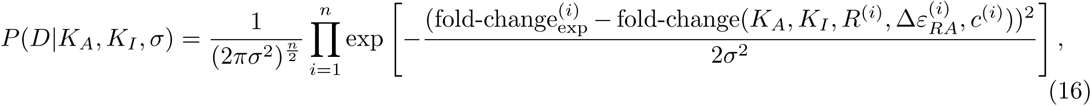

where fold-change 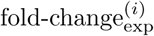 is the experimental fold-change and 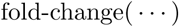 is the theoretical prediction. The product 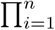 captures the assumption that the *n* data points are independent. Note that the likelihood and prior terms now include the extra unknown parameter *σ*. In applying Eq. (16), a choice of *K_A_* and *K_I_* that provides better agreement between theoretical fold-change predictions and experimental measurements will result in a more probable likelihood.

Both mathematically and numerically, it is convenient to define 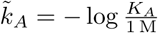 and 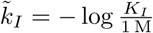 and fit for these parameters on a log scale. Dissociation constants are scale invariant, so that a change from 10 *μ*M to 1 *μ*M leads to an equivalent increase in affinity as a change from 1 *μ*M to 01 *μ*M. With these definitions we assume for the prior 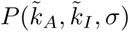 that all three parameters are independent. In addition, we assume a uniform distribution for 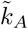 and 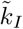 and a Jeffreys prior [36] for the scale parameter *σ*. This yields the complete prior

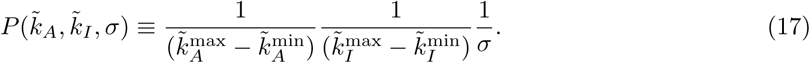

These priors are maximally uninformative meaning that they imply no prior knowledge of the parameter values. We defined the 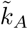 and 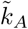 ranges uniform on the range of −7 to 7, although we note that this particular choice does not affect the outcome provided the chosen range is sufficiently wide.

Putting all these terms together we can now sample from 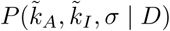 using Markov chain Monte Carlo (see GitHub repository) to compute the most likely parameter as well as the error bars (given by the 95% credible region) for *K_A_* and *K_I_*.

### Data Curation

All of the data used in this work as well as all relevant code can be found at this dedicated website. Data were collected, stored, and preserved using the Git version control software in combination with off-site storage and hosting website GitHub. Code used to generate all figures and complete all processing step as and analyses are available on the GitHub repository. Many analysis files are stored as instructive Jupyter Notebooks. The scientific community is invited to fork our repositories and open constructive issues on the GitHub repository.

## Acknowledgements

This work has been a wonderful exercise in scientific collaboration. We thank Hernan Garcia for information and advice for working with these bacterial strains, Pamela Björkman and Rachel Galimidi for access and training for use of the Miltenyi Biotec MACSQuant flow cytometer, and Colin deBakker of Milteny Biotec for useful advice and instruction in flow cytometry. The experimental front of this work began at the Physiology summer course at the Marine Biological Laboratory in Woods Hole, MA operated by the University of Chicago. We thank Simon Alamos, Nalin Ratnayeke, and Shane McInally for their work on the project during the course. We also thank Suzannah Beeler, Justin Bois, Robert Brewster, Ido Golding, Soichi Hirokawa, Jané Kondev, Tom Kuhlman, Heun Jin Lee, Muir Morrison, Nigel Orme, Alvaro Sanchez, and Julie Theriot for useful advice and discussion. We are also grateful to the three anonymous reviewers for substantially improving the quality of our work and our paper. This work was supported by La Fondation Pierre-Gilles de Gennes, the Rosen Center at Caltech, and the National Institutes of Health DP1 OD000217 (Director’s Pioneer Award), R01 GM085286, and 1R35 GM118043-01 (MIRA). Nathan Belliveau is a Howard Hughes Medical Institute International Student Research fellow.

## Competing interests

The authors have declared that no competing interests exist.

## Author contributions

MRM, SB, NB, GC, and TE contributed equally to this work. MRM, SB, NB, GC performed experiments. TE and MRM laid groundwork for the model. MRM, SB, NB, GC, and TE performed the data analysis. MRM, GC, NB, and SB wrote code used for all experimental analysis and parameter estimation. GC made the figures for the main text and GC, MRM, SB, and NB made figures for the supplemental information. MRM, SB, NB, GC, TE, and RP wrote the paper. ML and RP provided useful insight and advice in designing and executing the work.

